# *Structures of chloramphenicol acetyltransferase III and E. coli β-ketoacylsynthase III* co-crystallized with partially hydrolysed acetyl-oxa(dethia)CoA

**DOI:** 10.1101/2022.08.24.505111

**Authors:** Aaron B. Benjamin, Lee M. Stunkard, Jianheng Ling, Jaelen N. Nice, Jeremy R. Lohman

## Abstract

Acetyl-CoA is a reactive metabolite that non-productively hydrolyzes in a number of enzyme active sites on the crystallization time frame. In order to elucidate enzyme:acetyl-CoA interactions leading to catalysis, acetyl-CoA substrate analogs are needed. One possible analog for use in structural studies is acetyl-oxa(dethia)CoA (AcOCoA), where the thioester sulfur of CoA is replaced by an oxygen. Here we present structures of chloramphenicol acetyltransferase III (CATIII) and *E. coli* ketoacylsynthase III (FabH) from crystals grown in the presence of partially hydrolyzed AcOCoA and the respective nucleophile. Based on the structures, the behaviour of AcOCoA differs between the enzymes, with FabH reacting with AcOCoA and CATIII being unreactive. The structure of CATIII reveals insight into the catalytic mechanism, with one active site of the trimer having relatively clear electron density for AcOCoA and chloramphenicol, and the other active sites having weaker density for AcOCoA. One FabH structure has a hydrolyzed AcOCoA product oxa(dethia)CoA (OCoA) and the other FabH structure has an acyl-enzyme intermediate with OCoA. Together these structures provide preliminary insight into the use of AcOCoA for enzyme structure-function studies with different nucleophiles.

**Synopsis:** Stable analogs of acetyl-CoA are needed to support structure-function studies of acetyltransferase enzymes. We report structures of two enzymes in the presence of an acetyl-CoA analog where the thioester is replaced by an ester.

## 1. Introduction

Chloramphenicol acetyltransferase III (CATIII) and *E. coli* ketoacylsynthase III (ecFabH) both transfer an acetyl group from acetyl-CoA to an acceptor, a hydroxyl and thiol/thiolate, respectively, Figure 1. While these enzymes don’t share any sequence or structural conservation, they both display negative cooperativity in acetyl-CoA substrate binding and catalysis.(Ellis *et al*., 2002, Alhamadsheh *et al*., 2007) The function of negative cooperativity with respect to catalysis is speculative due to a lack of ternary complex structures. Thus, having structures of these enzymes in complex with substrates would provide insights into the fundamental enzyme-substrate and protein-protein interactions leading to catalysis and cooperativity. The inherent equilibrium of the CATIII and EcFabH reactions with acetyl-CoA lie so far toward the products that substrate bound states are difficult to determine. In addition, the inherent reactivity of the acetyl-CoA thioester and non-productive activation by the enzyme often leads to hydrolysis during crystallization. While there are new methods that may be useful for overcoming the inherent side reactivity of acetyl-CoA during co-crystallization, such as using free-electron lasers and serial crystallography, not every system will be amenable as the crystals must survive conformational changes linked to substrate binding.(Chapman, 2019) A tried-and-true method is the use of suitable stable substrate or transition state analogs.

**Figure 1.**
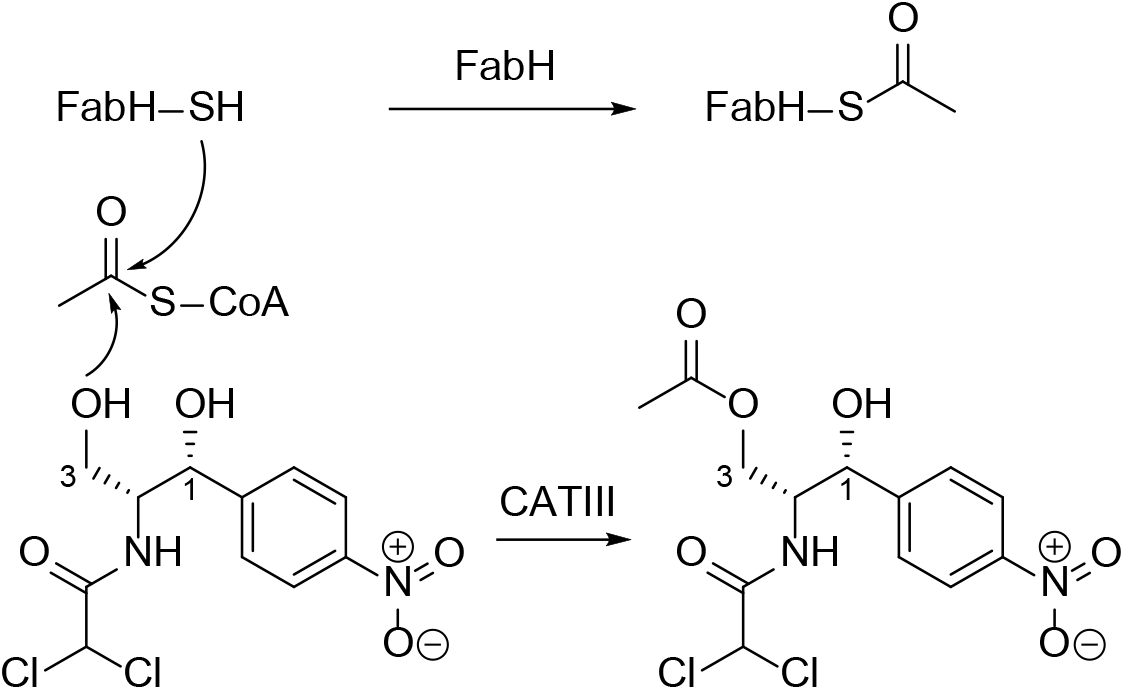
The transthiolation step of FabH and catalytic activity of CATIII.

A substrate analog of both CATIII and ecFabH is acetyl-oxa(dethia)CoA (AcOCoA), which is expected to be more stable than acetyl-CoA in structure-function studies.(Jencks & Gilchrist, 1964, Yang & Drueckhammer, 2001) We recently synthesized AcOCoA and similar analogs to support structure-function studies.(Stunkard, Kick, *et al*., 2021, Stunkard, Benjamin, *et al*., 2021, Stunkard *et al*., 2019) However, there was a report of the AcOCoA synthesis before ours.(Weeks *et al*., 2018) In that study, a similar substrate analog, fluoroacetyl-oxa(dethia)CoA was hydrolyzed 500-fold slower by a thioesterase than the native fluoroacetyl-CoA substrate. A truncated AcOCoA substrate acetyl-oxa(dethia)pantetheine-pivoyl has been crystallized with a thiolase, where it did not participate in the transthiolation or carbon-carbon bond forming reactions.(Merilainen *et al*., 2008) Thus, we initially expected AcOCoA to be relatively stable in CATIII and ecFabH crystals, both of which crystallize overnight. However, during stability assays we found that ecFabH was able to hydrolyze AcOCoA to oxa(dethia)CoA (OCoA), albeit extremely slowly compared to acetyl-CoA.(Boram *et al*., 2022) While improving on the synthesis of AcOCoA, we pursued crystal structures of the model enzymes CATIII and ecFabH with samples of AcOCoA that contained approximately 25-35% OCoA.

CATIII has long been a model enzyme for structure-function studies with relevance to antibiotic drug development.(Shaw & Leslie, 1991) With the recent structure of the ribosome with chloramphenicol bound, there should be renewed interest in finding analogs that retain ribosome inhibition and overcome CATIII and other chloramphenicol acetyltransferase activities.(Svetlov *et al*., 2019) Based on the binding of CoA and chloramphenicol, a hypothesis was developed where the chloramphenicol 1-hydroxyl stabilized a water that acts as a member of the oxyanion hole.(Lewendon & Shaw, 1993, Lewendon *et al*., 1990) Thus, some of the catalytic activity and substrate specificity might come from the chloramphenicol substrate itself, in other words, a substrate assisted catalysis hypothesis. The chloramphenicol analog lacking the 1-hydroxyl is a much poorer substrate, supporting the hypothesis. A ternary structure of AcOCoA and chloramphenicol bound to CATIII could confirm the Shaw hypothesis. Furthermore, CATIII is a trimer that displays negative cooperativity with respect to acetyl-CoA. A structure with AcOCoA could reveal conformational differences between the monomers leading to the cooperative behavior. Our structures here reveal a mixture of AcOCoA and OCoA bound, but enough to support the substrate assisted catalysis hypothesis. In addition, each active site of the biologically relevant trimer had varying levels of occupancy possibly revealing cooperativity.

In *E. coli* and other similar bacteria, FabH, carries out the first acyl carrier protein (ACP) dependent carbon-carbon bond forming step, making it a target for antibiotic drug development. The two active sites in the ecFabH homodimer appear to have negative cooperativity in the binding and/or reaction with acetyl-CoA.(Alhamadsheh *et al*., 2007) Once one FabH active site forms the acyl-enzyme intermediate, there is a large decrease in the race of acetylation of the second. Comparing crystal structures in the presence and absence of CoA reveals positional variations in loops that interact between the monomers in the dimer, which may explain some of the negative cooperativity. A convoluting factor is that the active site loops of FabH can be found in a disordered state (PDB 1HNK).(Qiu *et al*., 2001) The disordered state correlates with our recent finding that FabH displays significant hysteresis when presented with malonyl-CoA in the absence of acetyl-CoA as a substrate for decarboxylation.(Boram *et al*., 2022) Incubation with acetyl-CoA alleviates the hysteresis indicating FabH goes from a disordered state not competent for catalysis to one that is. However, the only structure with an acyl-enzyme intermediate bound has weak density that could be modeled as a partially bound acyl group and overlapping water (PDB 1HNH, electron density files not deposited).(Qiu *et al*., 2001) The ecFabH structures presented here with AcOCoA confirm that the acetyl group still gets transferred to generate the acyl-enzyme intermediate. These results reveal that other analogs such as acetyl-aza(dethia)CoA or acetyl-carba(dethia)CoA are needed to capture the acetyl-CoA substrate bound state of ecFabH.

The behaviors of CATIII and ecFabH with AcOCoA provides a comparison of how altering the electrophilic substrate from a thioester to an ester results in very different outcomes with respect to transition state stabilization. In one case, CATIII, the enzyme is unable to sufficiently stabilize the transition to the product, even on the crystallization time scale of days. Whereas with ecFabH, the enzyme is able to generate the thermodynamically unfavorable thioester, albeit with a large excess of substrate, within the same timeframe as CATIII. In order to fully comprehend how these enzymes carry out their reactions, it is likely neutron diffraction will be needed to confirm the positions of hydrogens, which are key for catalysis.

## 2. Materials and methods

### 2.1. Macromolecule production

Cloning and protein expression was done as previously reported for FabH.(Boram *et al*., 2022) Briefly, *fabH* from *E. coli* K-12 genomic DNA (UniProt P0A5R0) was cloned into a pRSF derived vector with a TEV protease cleavable site between the protein and N-terminal hexa-histidine tag. We found the addition of two glycines between the TEV site and FabH N-terminus were necessary for efficient tag cleavage. The CATIII gene was synthesized by Integrated DNA Technologies according to the sequence found in the transmissible plasmid R387 (UniProt P00484) with appropriate overhangs for Gibson cloning into the modified pRSF vector. Again, double glycines were added to facilitate efficient TEV protease cleavage, CATIII+GG. For CATIII+GG, the primers used were (additional codons underlined yielding N-terminal sequence <u>SGG</u>NYTK….):

CATIII+GG-forward 5′-GAGAACCTCTACTTCCAA*<u>AGTGGTGGT</u>AACTATACAAAATTTGATG*-3′,

CATIII+GG-reverse 5′-CTCGAGGAGATTACGGA*TTATTTTAATTTACTGTTACAC*-3′,

Plasmids with *fabH*+GG, and *catIII*+GG were transformed into *E. coli* BL21 (DE3) used as the expression strain. Overnight cultures were used to inoculate LB containing 10 mM MgCl_2_, trace metals, and 50 µg/mL kanamycin, which were incubated at 37 °C and shaken at 180 rpm. Upon reaching an OD_600_ of ∼0.5-0.6, the temperature was reduced to 18 °C. Once the cultures reached thermal equilibrium, gene expression was induced by the addition of isopropyl β-d-thiogalactopyranoside with a final concentration of 500 µg/ml, with incubation for an additional 16 hours. *E. coli* cells were harvested by centrifugation at 6300 rpm and 4 °C for 30 min.

*E. coli* cell pellets, carrying FabG+GG or CATIII+GG were re-suspended in lysis buffer (1 μg/ml DNase, 1 μg/ml lysozyme, 300 mM NaCl, 20 mM imidazole, 10% glycerol, and 20 mM Tris-HCl pH 8.0), sonicated (60 × 1 s on ice), and clarified by centrifugation at 20000 g and 4 °C for 30 min. The supernatant was filtered, applied to a 5 ml HisTrap HP (GE Healthcare), and washed with lysis buffer using an Äkta pure fast-performance liquid chromatography system (GE Healthcare,). Wash buffer (300 mM NaCl, 40 mM imidazole and 20 mM Tris-HCl pH 8.0) was used to remove additional contaminants, and proteins were eluted with wash buffer containing 500 mM imidazole. At this point the purity of FabG+GG and CATIII+GG from the fractions was analyzed by sodium dodecyl sulfate– polyacrylamide gel electrophoresis (SDS–PAGE). Pure fractions were pooled and cleaved using TEV protease to remove the 6His tag. FabG+GG and CATIII+GG were then buffer-exchanged into storage buffer (200 mM NaCl, 10 mM Tris-HCl pH 8.0), concentrated, and frozen in small aliquots with liquid nitrogen, and stored at -80 °C.

### 2.2. Crystallization

FabG+GG and CATIII+GG were screened against 384 crystallization conditions in 500 nL sitting drops at 20°C, set up with a Mosquito (TTPlabtech, Melbourne, Australia) to find initial conditions with AcOCoA (partially hydrolysed). FabH+GG at 21 mg/mL with 10 mM AcOCoA produced crystals by the hanging drop method over 1.0 mL wells containing 1.5% DMSO, 23% PEG 3350, 75 mM MgCl_2_, and 100 mM HEPES-NaOH pH 7.5 or 20% PEG 6000, 3% PEG 400, and 100 mM MgCl_2_. CATIII+GG at 24 mg/mL with 10 mM AcOCoA and saturating concentrations of chloramphenicol (solid chloramphenicol was added to the protein and AcOCoA until powder remained in solution followed by centrifugation to precipitate excess material) produced crystals by the hanging drop method over 1.0 mL wells containing 46% PEG 400, 100 mM MgCl_2_, 0.1 M sodium citrate pH 5.5, in 4 µL drops (1:3, protein:well). Crystals were looped and frozen directly out of the drops with liquid nitrogen.

### 2.3. Data collection and processing

X-ray diffraction data for all datasets were collected at Advanced Photon Source LS-CAT beamline 21-ID-G at a wavelength of 0.97856. Diffraction intensities were integrated, reduced, and scaled using HKL2000,(Otwinowski & Minor, 1997) with data collection and refinement statistics listed in Tables 3 and 4.

### 2.4. Structure solution and refinement

Molecular replacement with the program Phaser was used to solve our structures of FabH with OCoA (PDB: 6×7R) or OCoA and acetylated-FabH (PDB: 6×7S) based off PDB 1HNJ coordinates(Qiu *et al*., 2001), and CATIII with AcOCoA (PDB: 6×7Q) was solved based off PDB 3CLA coordinates.(Leslie, 1990) Refinement was conducted using Refmac (Murshudov *et al*., 1997) in the CCP4i package (Winn *et al*., 2011) with automated model building performed with ARP/wARP (Langer *et al*., 2008) and manual model building with Coot.(Emsley *et al*., 2010)

## 3. Results and discussion

### 3.1. Structure of CATIII in complex with AcOCoA and chloramphenicol

We co-crystallized CATIII in the presence of our partially hydrolyzed AcOCoA and chloramphenicol. This produced crystals that grew overnight in various conditions. The vast majority of crystals gave diffraction patters that were difficult to index due to what appeared to be twinning problems. Serendipitously, we found a single large crystal that diffracted well, was easily indexed in the primitive tetragonal Bravis lattice and solved in the P4_2_2_1_2 spacegroup with a trimer in the asymmetric unit. The final structure had good refinement statistics and all residues for each monomer could be modeled. Although no transition metals were added to the crystallization conditions, we found positions for two metals that we modeled as Zn^2+^ based on the CheckMyMetal validation server, which would have come from the inclusion of trace metals in the expression media.(Zheng *et al*., 2017) Alternatively, these metals might be Ni^2+^ that leached from the Ni-NTA column during purification. The metals are liganded by Glu18 and His22 in each chain, and the same residues in a symmetry mate trimer, with the pattern: chain A interacts with symmetry chain C, and chain B interacts with symmetry chain B, creating a larger order hexamer. The previous deposited structures of CATIII were in spacegroup R32, with a single molecule per asymmetric unit (we would like to note the numbering scheme of Leslie and Shaw added 5 residues to 1-74 and 6 residues to 75-213, here we use the linear numbering of CATIII).(Leslie, 1990, Leslie *et al*., 1988) The R32 CATIII structures had two Co^2+^ metals bound (0.5 mM added to crystallization condition), one site is shared with a metal in our structure that makes the hexamer, while the other Co^2+^ occupies a special position situated between the backbone carbonyls of Asn63 and Asp81 with rather long interaction distances, ∼4 Å. While the shared metals generate a larger order hexamer in both crystal forms, all other crystal contacts are essentially unique. Differences in the N-termini of the CATIII from previous studies and ours likely lead to the crystal packing variations. The previous R32 structures have an N-terminus starting at Met1 and the N-terminal amine has good packing with a hydrogen bond to the carbonyl of Lys211 that likely stabilizes the crystals and isn’t possible without a free amine. Our construct has an N-terminus with a Ser-Gly-Gly-Asn2 cloning artifact that would disrupt the packing seen in the R32 structures. The R32 crystal packing was reported to deteriorate upon exposure to CoA, preventing elucidation of acetyl-CoA or analog binding via soaking.(Leslie *et al*., 1986) Thus, our structure provides insight into engineering crystal contacts that allow production of trimers in the asymmetric unit in order to understand the subtle conformational changes associated with negative cooperativity.

Our CATIII structure has clear electron density for chloramphenicol in each monomer, which resides in exactly the same orientation as in previous structures. The electron density for the AcOCoA/OCoA is relatively clear in chain A, somewhat clear in chain B and difficult to model in chain C, Figure 2. We take the differences in electron density to correspond to negative cooperativity, but differences in crystal packing can’t be ruled out. The AcOCoA in chain A participates in crystal packing and is not free to leave, while AcOCoA in chains B and C are open to solvent. A comparison of the chains shows slight differences in the loops surrounding the CoA binding pocket in chain C, further supporting the idea of our electron density reflecting negative cooperativity.

**Figure 2.**
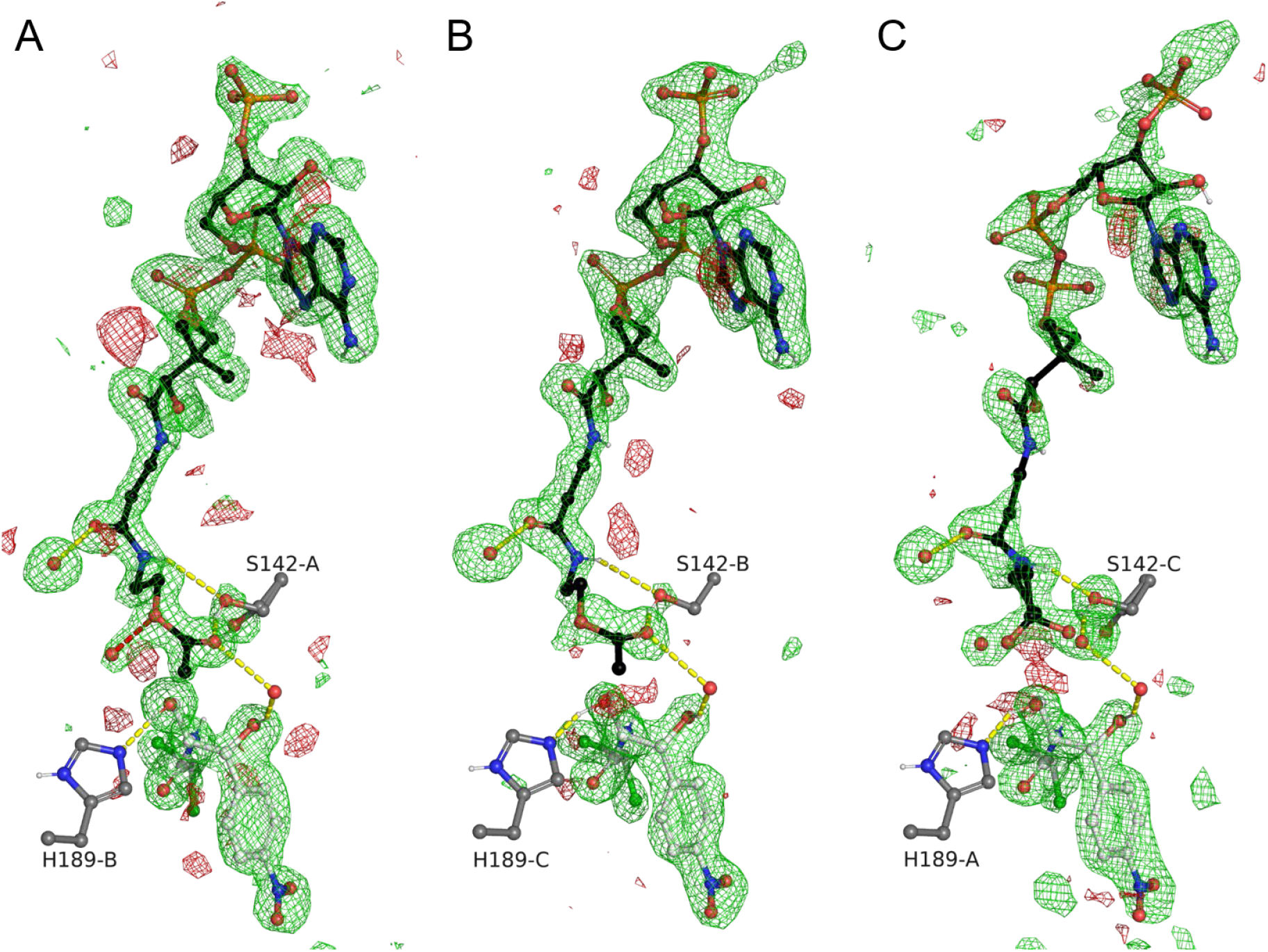
Electron density for CATIII ligands in the trimer active sites. AcOCoA or OCoA shown in black sticks, chloramphenicol shown in white sticks and labelled CATIII active site residues in gray sticks. The σA-weighted mF_o_-DF_c_ maps for omitted ligands shown at +3σ in green and -3σ in red as 5Å bricks. The Ser142 side-chain hydroxyl was also omitted in order to judge occupancy. Notice a close water molecule to the ester oxygen in A) lies in a similar position as a partially occupied water in C), revealing some presence of OCoA in A).

In monomer A, we can clearly model the position of the acetyl group, even though the electron density supports about 20% OCoA, in which case a water molecule takes the place of the acetyl ketone oxygen, Figure 2. The binding of the acetyl-group is pretty much exactly what Leslie and Shaw predicted based on a structure with coenzyme A bound (structure not deposited in the PDB), Figure 3.(Shaw & Leslie, 1991, Leslie *et al*., 1988) The location of a water bound between the acetyl ketone and chloramphenicol 1-hydroxyl suggests that it stabilizes the transition states in concert with His189. This is the first time a substrate analog of acetyl-CoA has been solved in the CATIII active site, revealing an active site water interacts with the tetrahedral intermediate, confirming the substrate assisted catalysis hypothesis. Nevertheless, the positions of the hydrogens that are key for catalysis remain to be determined. Our preliminary structures here provide a platform upon which to perform neutron diffraction to obtain a clear idea of how the transition state is set up.

**Figure 3.**
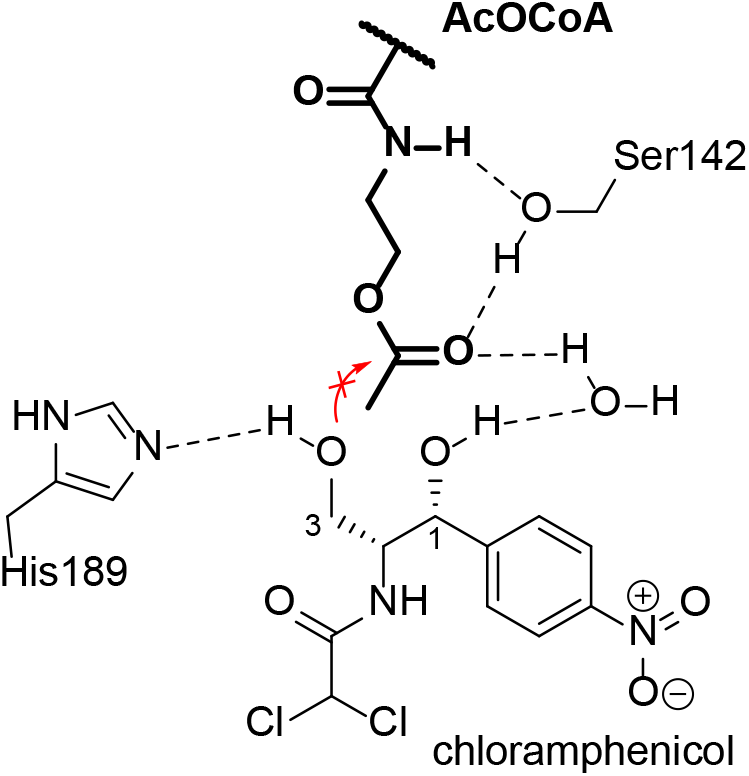
Active site interactions of AcOCoA with CATIII and chloramphenicol co-substrate. Notice a water bound to chloramphenicol helps create the oxyanion hole.

### 3.2. Structures of FabH in complex with OCoA

We co-crystallized ecFabH in the presence of our partially hydrolyzed AcOCoA. This produced crystals that grew overnight in various conditions. Crystals grown from PEG and Mg^2+^ diffracted well and were indexed in the primitive tetragonal Bravis lattice and solved in the P4_1_2_1_2 spacegroup (a = b = 73 Å) each with a monomer in the asymmetric unit, Table 1. The final structures had good refinement statistics and all residues for each monomer could be modeled. The previous structures of ecFabH fall into one of three crystal forms, crystal type I is identical to ours reported here while the others are primitive orthrombic. Crystal type II is P2_1_2_1_2_1_, a = ∼63 Å, b = ∼65 Å, c = ∼163 Å, and crystal type III is P2_1_2_1_2_1_, a = ∼64 Å, b = ∼81 Å, c = ∼122 Å, with both having a dimer in the asymmetric unit. Solid density for the CoA (PDB 1HND, 1HNJ) and partial density for the acyl enzyme intermediate (PDB 1HNH) have only been seen in crystal type I,(Qiu *et al*., 2001) with weak CoA density in crystal type II (PDB 1EBL)(Gajiwala *et al*., 2009) and crystal type III (PDB 2EFT and 2GYO).(Alhamadsheh *et al*., 2007) Based on structural alignments between crystals type I and II/III, it appears that an N-terminal his tag may be responsible for favoring type II/III crystals, due to clashes that would be present in type I. In our crystals, we have some disorder in the Ser-Gly-Gly-Met1 tag artifact; however, there is enough room that the tag artifact still allows tight crystal packing found in crystal form I. Elongating the tag may be a way to favor production of the type II/III crystals, which would be helpful for capturing differences between the active sites associated with negative cooperativity.

**Table 1.**
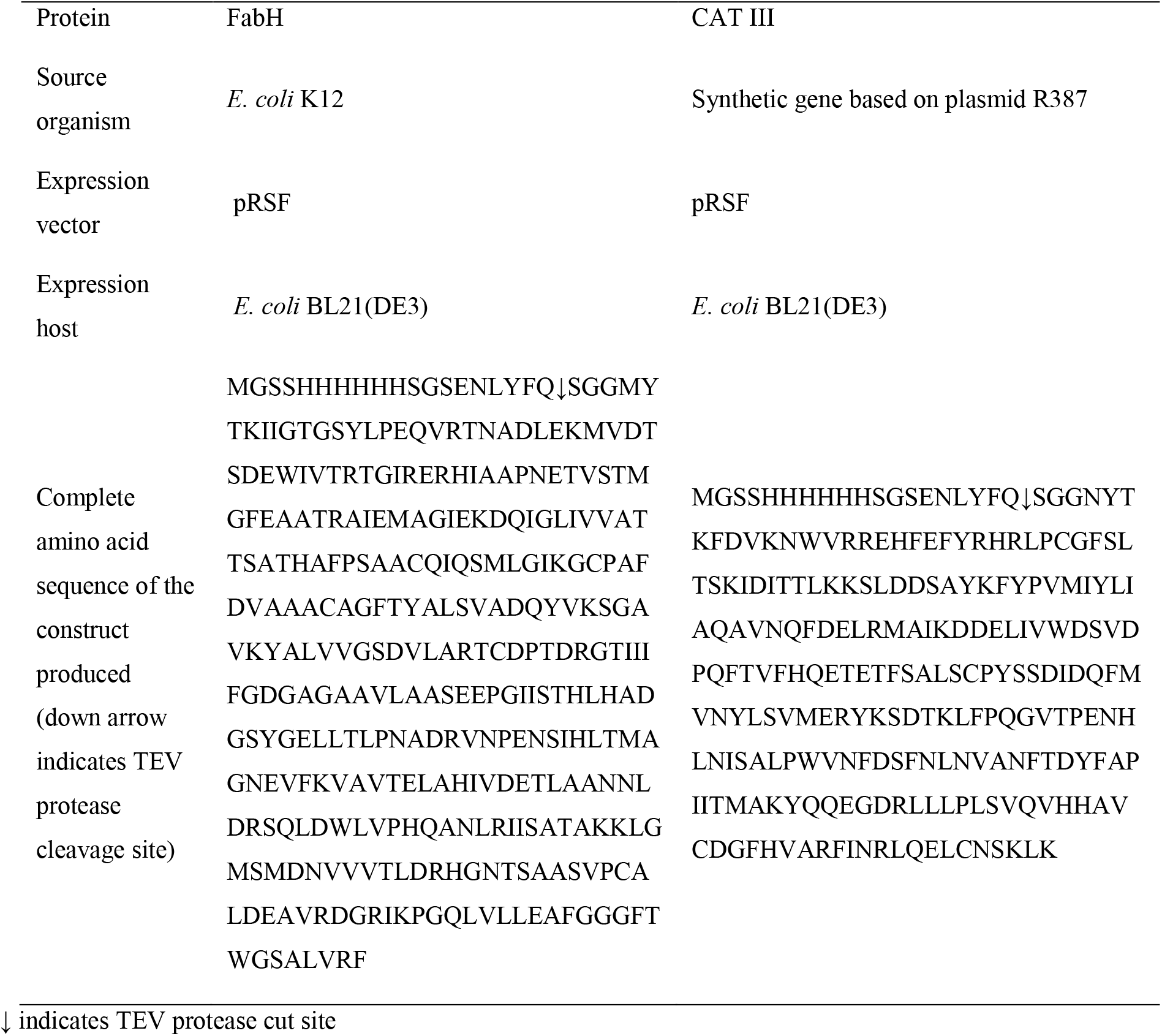
Macromolecule production information

**Table 2.** Crystallization

|  | Ac-FabH+OCoA | FabH + OCoA | CATIII + AcOCoA |
| --- | --- | --- | --- |
| Method | Vapor diffusion, hanging drop | Vapor diffusion, hanging drop | Vapor diffusion, hanging drop |
| Plate type | VDX | VDX | VDX |
| Temperature (K) | 298 | 298 | 298 |
| Protein concentration | 680 μM | 680 μM | 960 μM |
| Buffer composition of protein solution | 10 mM AcOCoA, 200 mM NaCl, 10 mM Tris-HCl pH 8.0 | 10 mM AcOCoA, 200 mM NaCl, 10 mM Tris-HCl pH 8.0 | 10 mM AcOCoA, saturated chloramphenicol, 200 mM NaCl, 10 mM Tris-HCl pH 8.0 |
| Composition of reservoir solution | 75 mM MgCl<sub>2</sub>, 20% PEG6000, 3% PEG400 | 1.5% DMSO, 75 mM MgCl<sub>2</sub>, 23% PEG3350, 100 mM HEPES-NaOH pH 7.5 | 100 mM MgCl<sub>2</sub>, 46% PEG400, 100 mM sodium citrate pH 5.5 |
| Volume and ratio of drop | 4 μl 2:2 protein:well | 4 μl 2:2 protein:well | 4 μl 1:3 protein:well |
| Volume of reservoir | 1.0 mL | 1.0 mL | 1.0 ml |

**Table 3.** Data collection and processing Values for the outer shell are given in parentheses.

|  | Ac-FabH + OCoA | FabH + OCoA | CATIII + AcOCoA |
| --- | --- | --- | --- |
| Diffraction source | APS beamline 21-ID-G | APS beamline 21-ID-G | APS beamline 21-ID-G |
| Wavelength (Å) | 0.97856 | 0.97856 | 0.97856 |
| Temperature (K) | 80 | 80 | 80 |
| Space group | *P4<sub>1</sub>2<sub>1</sub>2* | *P4<sub>1</sub>2<sub>1</sub>2* | *P4<sub>2</sub>2<sub>1</sub>2* |
| *a*, *b*, *c* (Å) | 72.63, 72.63, 102.87 | 72.63, 72.63, 102.87 | 106.92, 106.92, 126.60 |
| $\alpha$, $\beta$, $\gamma$ (°) | 90, 90, 90 | 90, 90, 90 | 90, 90, 90 |
| Resolution range (Å) | 50–1.30 (1.35–1.30) | 50.0–1.35 (1.40–1.35) | 50.0–1.68 (1.74–1.68) |
| Total No. of reflections | 247744 | 224532 | 311296 |
| No. of unique reflections | 68034 | 60608 | 84122 |
| Completeness (%) | 99.6 (96.5) | 98.7 (100) | 99.8 (100) |
| Redundancy | 14.2 (12.0) | 13.9 (13.7) | 14.5 (14.3) |
| $\langle I/\sigma(I) \rangle$ | 34.4 (1.94) | 25.1 (3.85) | 23.5 (2.97) |
| *R*<sub>r.i.m.</sub> | 0.070 (0.568) | 0.089 (0.797) | 0.152 (1.308) |
| *R*<sub>p.i.m.</sub> | 0.019 | 0.024 | 0.040 |
| Overall *B* factor from Wilson plot (Å<sup>2</sup>) | 14.74 | 20.81 | 20.37 |

**Table 4.**
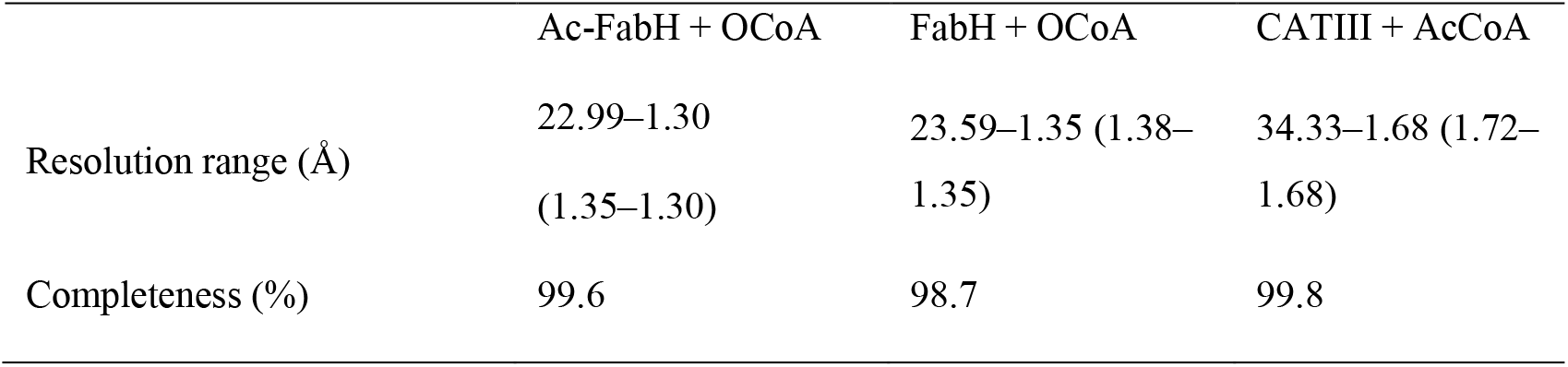
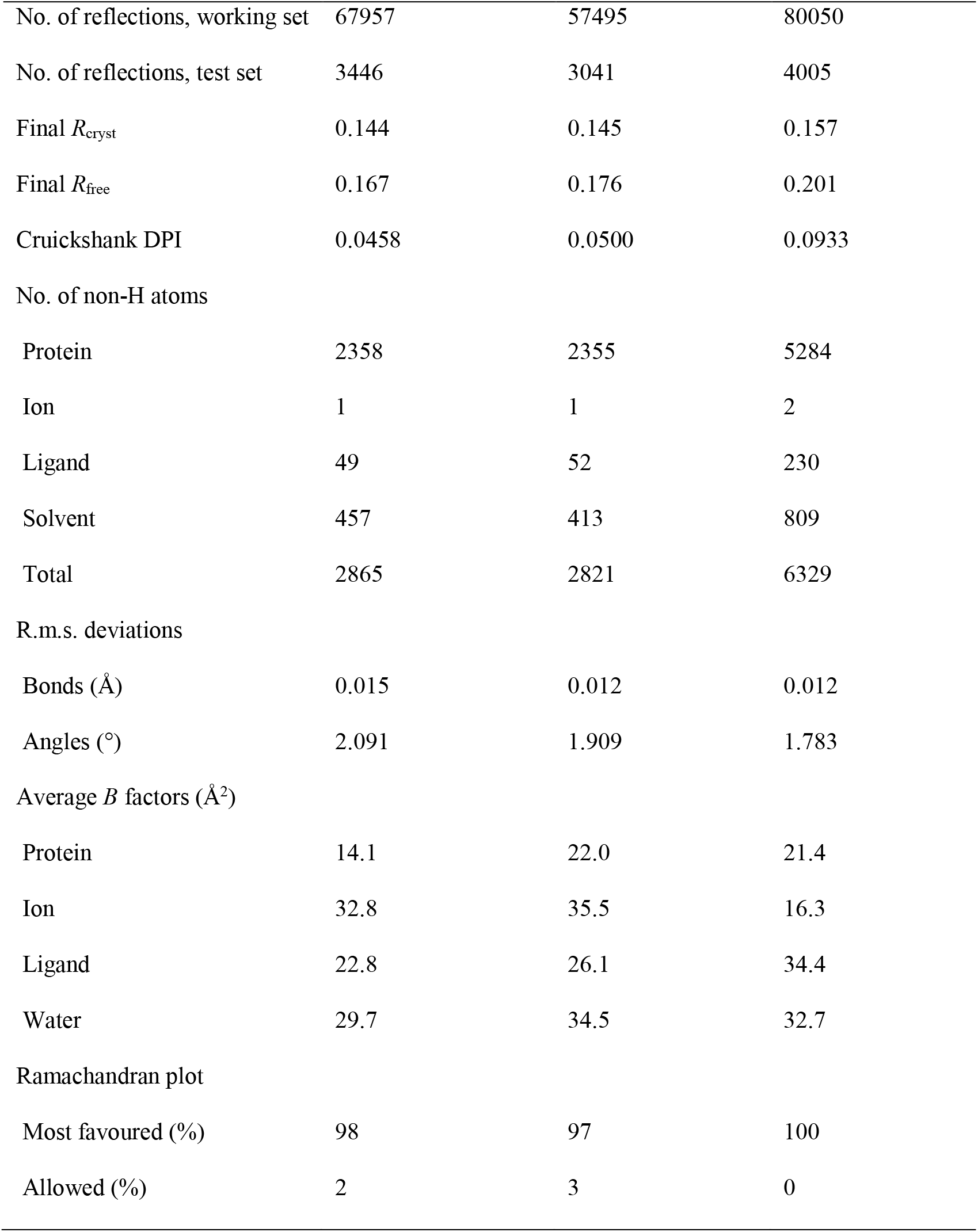
Structure solution and refinement Values for the outer shell are given in parentheses.

|  | Ac-FabH + OCoA | FabH + OCoA | CATIII + AcCoA |
| --- | --- | --- | --- |
| Resolution range (Å) | 22.99–1.30<br>(1.35–1.30) | 23.59–1.35 (1.38–<br>1.35) | 34.33–1.68 (1.72–<br>1.68) |
| Completeness (%) | 99.6 | 98.7 | 99.8 |
| No. of reflections, working set | 67957 | 57495 | 80050 |
| No. of reflections, test set | 3446 | 3041 | 4005 |
| Final $R_{\text{cryst}}$ | 0.144 | 0.145 | 0.157 |
| Final $R_{\text{free}}$ | 0.167 | 0.176 | 0.201 |
| Cruickshank DPI | 0.0458 | 0.0500 | 0.0933 |
| No. of non-H atoms |  |  |  |
| Protein | 2358 | 2355 | 5284 |
| Ion | 1 | 1 | 2 |
| Ligand | 49 | 52 | 230 |
| Solvent | 457 | 413 | 809 |
| Total | 2865 | 2821 | 6329 |
| R.m.s. deviations |  |  |  |
| Bonds (Å) | 0.015 | 0.012 | 0.012 |
| Angles (°) | 2.091 | 1.909 | 1.783 |
| Average $B$ factors (Å<sup>2</sup>) | | | |
| Protein | 14.1 | 22.0 | 21.4 |
| Ion | 32.8 | 35.5 | 16.3 |
| Ligand | 22.8 | 26.1 | 34.4 |
| Water | 29.7 | 34.5 | 32.7 |
| Ramachandran plot |  |  |  |
| Most favoured (%) | 98 | 97 | 100 |
| Allowed (%) | 2 | 3 | 0 |

Slight differences in the crystallization conditions for our structures resulted in differing amounts of density for the acyl-enzyme intermediate and OCoA product, Figure 4. The B-factors for the carbons of the acyl-enzyme intermediate acetyl-group are ∼1.5-2 times as large as the atoms of the cysteine to which they are attached, suggesting maybe 50% occupancy. Our structure with electron density for the acyl-enzyme intermediate has relatively poor electron density for the bound OCoA. Our structure with excellent density for OCoA has no density for the acyl-enzyme intermediate. Taken together, we can get a clear view of how the enzyme interacts with the products of AcOCoA. The acyl-enzyme intermediate is very similar in structure to that previously determined with acetyl-CoA (PDB 1HNH).(Qiu *et al*., 2001) Similarly, our structure with clear OCoA bound, is almost identical to having CoA bound (PDB IHND, 1HNJ).

**Figure 4.**
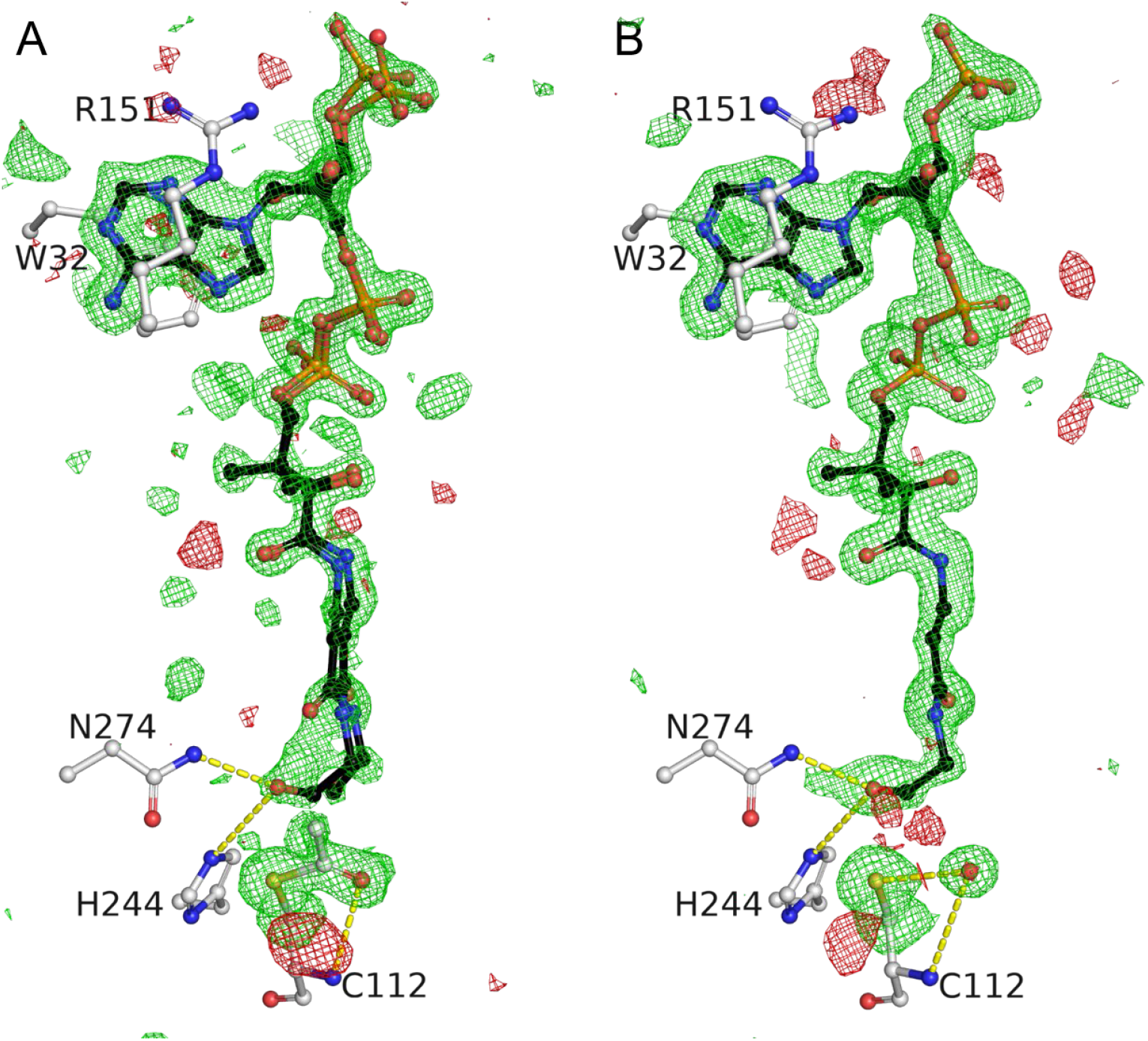
Electron density for FabH ligands. OCoA is shown in black sticks and labelled FabH active site residues in white sticks. The σA-weighted mF_o_-DF_c_ maps shown at +3σ in green and -3σ in red as 5Å bricks for A) omitted OCoA and Cys112 Cβ and sulphur or B) Cys112 Cβ, sulphur and acetyl-group. A) is Ac-FabH + OCoA and B) is FabH + OCoA. Notice a water molecule takes the place of the acetyl-cysteine carbonyl.

Unfortunately, due to the presence of only a monomer in our asymmetric unit, it is difficult to gain insight into the negative cooperativity displayed by the enzyme. Nevertheless, careful inspection of the electron density maps, and comparisons with other structures reveals conformational heterogeneity that might help explain some of the cooperative behavior. The α-helix between Leu249-L258 has spurious density, suggesting the helix is in an alternate conformation part of the time. Alignment of our structures with structures 3IL9 (Gajiwala *et al*., 2009) or 1EBL (Davies *et al*., 2000) reveals the secondary conformation of this loop, likely reflecting a state without CoA bound, Figure 5. Questions remain concerning the transthiolation reaction; is the active site cysteine in the thiol or thiolate state, and how the protonation state of His244 changes before and after formation of the acylenzyme intermediate. The multiple water and nearby side chain conformations suggest we don’t have the data to accurately interpret how the acyl-enzyme intermediate alters these protonation states. We expect that substrate analogs such as acetyl-aza(dethia)CoA or acetyl-carba(dethia)CoA are needed to capture a “substrate bound” state of ecFabH and to overcome the conformational heterogeneity found in these FabH structures.

**Figure 5.**
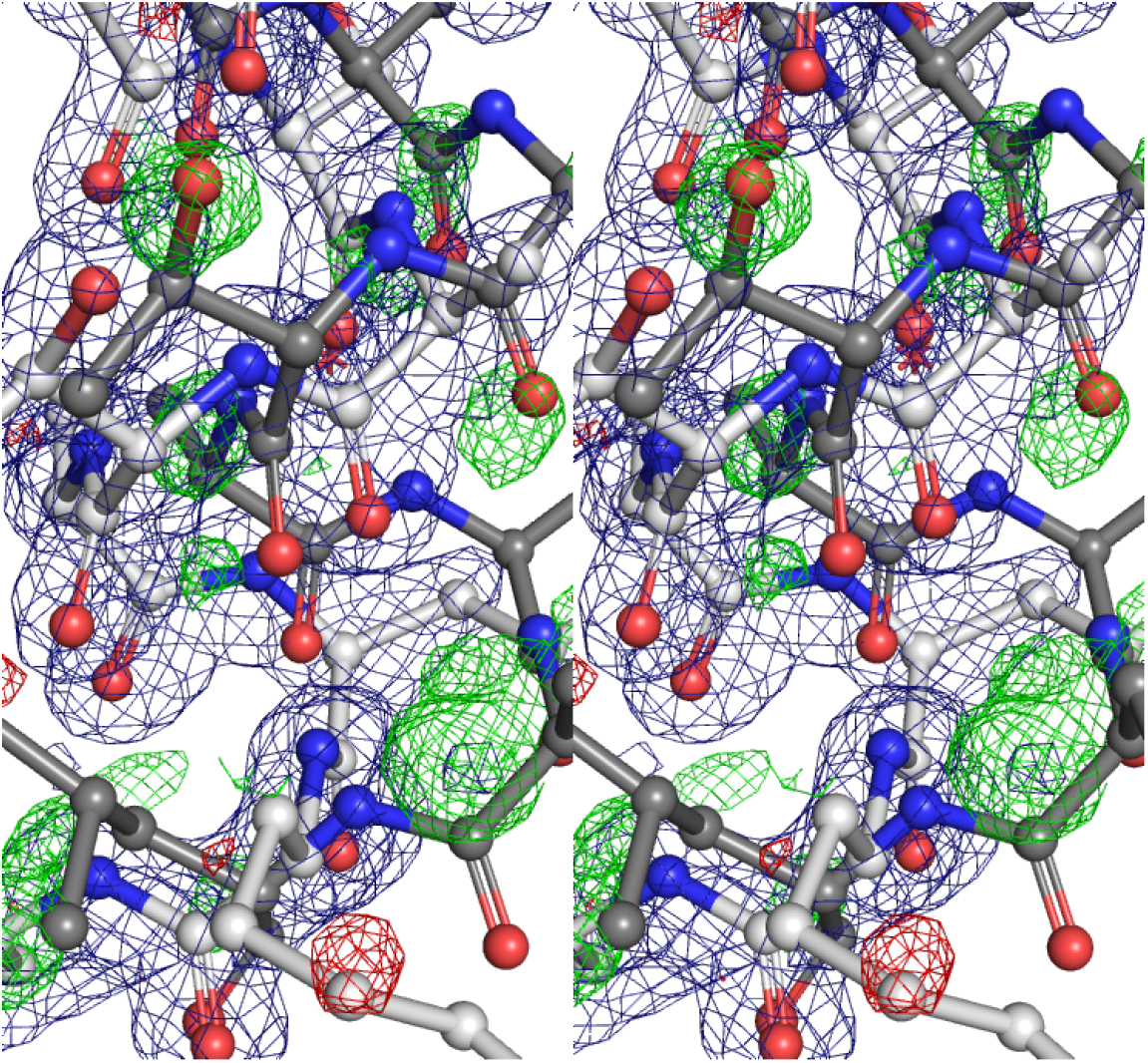
FabH alternative conformation for residues 249-258 shown as a stereoview. The σA-weighted 2mF_o_-DF_c_ map is shown in blue and mF_o_-DF_c_ maps shown at +3σ in green and -3σ in red for the Ac-FabH + OCoA structure. The Ac-FabH + OCoA is shown in white sticks, and the residues from PDB 1EBL chain A are shown in gray. Notice how the residues for 1EBL occupy the residual positive density suggesting some fraction is populated in our crystals.

### 3.3. Different behaviors for AcOCoA in CATIII/FabH active sites

CATIII does not perform the forward reaction between AcOCoA and chloramphenicol during the crystallization time frame at pH 5.5. The ΔG for the reaction is expected to be 0, which should give us an equal amount of substrate and product. In our case we used a large excess of chloramphenicol, which should have driven the reaction forward leaving only OCoA. There are a couple explanations for the lack of acyl-transfer in this context. The ΔG^‡^ is inherently too large for transesterification to be efficiently overcome by the enzyme due to geometric or electronic properties of the ester compared to the thioester. The crystallographic pH is far enough below the pK_a_ of the active site histidine (6.3) inhibiting the reaction.(Shaw & Leslie, 1991) We expect follow up studies examining the reaction at various pH can shed light on which of these two hypotheses is correct.

FabH does hydrolyze AcOCoA very slowly under relatively dilute conditions used in enzymology experiments, compared to the crystallographic conditions used here where AcOCoA is only in ∼10 fold excess, as such the rate of FabH hydrolysis is a concern.(Boram *et al*., 2022) Our structural studies here confirm that the hydrolysis can occur through the acyl-enzyme intermediate. The formation of the acyl-enzyme intermediate is somewhat unexpected, as it is unfavorable with a positive ΔG. We used ∼10-fold more AcOCoA than FabH in our crystallization condition, which may have contributed to the spontaneous formation of the acyl-enzyme intermediate. We observed a similar rate of hydrolysis for AcOCoA and acetyl-CoA in the absence of a malonyl-thioester substrate at pH 8, which is ten times higher than background hydrolysis in buffer, suggesting FabH does have a noticeable effect.(Boram *et al*., 2022) However, it did not appear that AcOCoA acted as an acyl donor, since the reaction in the presence of malonyl-CoA did not yield more product than malonyl-CoA alone. Similar to CATIII, it appears that geometric or electronic considerations are what become rate limiting in the reaction of FabH with AcOCoA. Future studies examining substrate binding of acetyl-CoA will require a different strategy than using AcOCoA such as more stable analogs.

Our structures demonstrate different behaviors for AcOCoA depending on the context. AcOCoA is a substrate analog for CATIII but a very slow substrate for FabH. It may be the case that hydroxyl acceptors in general will lead to substrate analog behavior and thiol acceptors will lead to product formation. With the aforementioned fluoroacetyl-CoA hydrolase, which generates a threonine based acyl-enzyme intermediate, fluoroacetyl-oxy(dethia)CoA was a substrate with a 500-fold slower rate.(Dias *et al*., 2010, Weeks *et al*., 2018, Weeks *et al*., 2010) Together it suggests that acyloxy(dethia)CoAs are likely to be useful tools for studying many enzymes with hydroxyl nucleophiles. The catalytic interactions for many Gcn5-related N-acetyltrasnferases (GNATs) still remains uncharacterized due to the spontaneous hydrolysis of acetyl-CoA and reactivity with their substrates. Our studies here with CATIII suggest that acyl-oxy(dethia)CoAs might be effective for capturing the ternary complexes; however, the reactivity with amine nucleophiles remains to be examined.

## Acknowledgements

This work was supported in part by NIH grant GM140290.

